# Warm and wet: robust lipase-producing bacteria from the indoor environment

**DOI:** 10.1101/148148

**Authors:** Kristie Tanner, Christian Abendrotht, Manuel Porcar

**Author notes:** Both authors contributed equally to this work (Kristie Tanner, Christian Abendrotht). Corresponding author: (MP).

## Abstract

Lipases are key biocatalysts with important biotechnological applications. With the aim of isolating robust lipolytic microbial strains, we have analyzed the bacterial communities inhabiting two domestic extreme environments: a thermophilic sauna and a dishwasher filter. Scanning electron microscopy revealed biofilm-forming and scattered microorganisms in the sauna and dishwasher sample, respectively. A culture-independent approach based on 16S rRNA analysis indicated a high abundance of Proteobacteria in the sauna sample; and, a large amount of Proteobacteria, Firmicutes, Cyanobacteria and Actinobacteria in the dishwasher filter. With a culture-dependent approach, we isolated 48 bacterial strains, screened their lipolytic activities on media with tributyrin as the main carbon source, and finally selected five isolates for further characterization. These strains, all of them identified as members of the genus *Bacillus,* displayed optimum lipolytic peaks at pH 6.5 and with 1-2% NaCl, and the activity proved very robust at a wide range of pH (up to 11.5) and added NaCl concentrations (up to 4%). The thermal, pH and salt robustness of the selected isolates is a valuable attribute for these strains, which are promising as highly tolerant biodetergents. To our knowledge, this is the first report regarding the isolation from an indoor environment of *Bacillus* strains with a high potential for industry.

## Introduction

In the past decade, research programs on indoor environments have resulted in an increasing data matrix of taxonomic and ecological interest [1, 2]. Attention has especially been paid to frequently used domestic places that are, on many occasions, overgrown with potential pathogenic bacteria, like in the recently described coffee-machine or refrigerator bacteriomes [3, 4]. It is important to stress that indoor environments mimic natural, often extreme, environments. For example, refrigerators are almost as cold as tundra and thus rich in cold-adapted bacteria, whereas sun-exposed artificial flat surfaces, such as solar panels, are home of a rich desert-like biocenosis [5]. Therefore, bioprospecting nearby indoor extreme environments is a poorly explored but yet promising screening strategy that might yield bacterial strains with new or improved biotechnological applications.

Indeed, and besides the obvious medical implications, another reason to further investigate indoor microbiomes is the search of enzymes with high industrial significance, especially as novel biocatalysts [6]. A very well known (natural) precedent is the discovery of the extremophile bacterium *Thermus aquaticus* [7], whose thermoresistant Taq polymerase allowed the revolutionary development of Polymerase Chain Reaction in the last decades of the 20th century.

Within the current repertoire of available enzymes, esterases are particularly suitable for industrial processes, since they are stable in organic solvents and can freely reverse the enzymatic reaction from hydrolysis to synthesis [8]. Lipases have also been highlighted as key biocatalysts for biotechnological applications, such as the production of new biopolymeric materials and biodiesel, or the synthesis of fine chemicals like therapeutics, agrochemicals, cosmetics and flavors [9]. Their stereoselective properties make them able to recognize enantiomers and enantiotopic groups, while many other enzymes for hydrolysis are just capable of metabolizing one antipode of the specific substrate [10].

Environments with extreme and/or oscillating temperatures are of special interest, due to the opportunity of finding esterases that are active at wide intervals of temperature and that can thus be used under a range of industrial conditions, such as those present in dishwashers or washing machines. A new and promising esterase has recently been discovered in the thermophilic bacterium *Thermogutta terrifontis.* This enzyme retains up to 95% of its activity after incubation for 1h at 80°C [11]. A cold-active and solvent-tolerant lipase from *Stenotrophomonas maltophilia* has also been reported, with retention of 57% of its activity at 5°C and more than 50% of its activity in pure organic solvents [12]. More examples of extremophilic enzymes with industrial potential include thermoalkalophilic esterases from *Geobacillus* sp., which have all proven active at high temperature (65°C) and at pH of up to 10 [13]; or a cold-adapted esterase from *Pseudoalteromonas arctica*, which still retained 50% of its activity at the freezing point of water [14].

Upon discovery, extremophile enzymes can often be further optimized to improve their industrial use, as it was the case for the thermal stability and activity in the cold-adapted lipase B from *Candida antarctica* through chemical linking of amino groups of the lipase to oxidized polysaccharides using reducing agents [15].

Bioprospecting indoor extreme environments could yield new lipolytic microbial strains harbouring previously uncharacterized esterases and other enzymes. In the present work, we have focused on the microbial communities inhabiting two high-temperature, domestic environments: a thermophilic sauna and a dishwasher. We have isolated 48 bacterial strains, many of them lipase-producing bacteria. Furthermore, we have characterized five of them, displaying robust lipase activities with promising biotechnological applications.

## Material and Methods

### 2.1. Sampling

Environmental samples were taken from a sauna and from a dishwasher. The sauna, set at a temperature of approximately 45°C and with 100% relative humidity, is a publicly-owned facility located in a communal swimming pool in Valencia (Spain) and therefore did not require specific permission for the sampling. A biofilm-like mass below the aluminium bench of the sauna was collected in a sterile 50 mL Falcon tube and was stored at -20 °C until required. The dishwasher sample was collected from the filter of a domestic Siemens dishwasher (property of one of the co-authors of this work, MP), Model sm6p1s. The sample was obtained by scratching the inner surface of the filter with a sterile bladder and the resulting biomass was kept at -20 °C until required.

### 2.2. Scanning electron microscopy

Biomass samples were fixed on a 0.2 μm membrane filter (Merck Millipore Ltd, Tullagreen, Cork, Ireland) using paraformaldehyde 2 % - glutaraldehyde 2.5 %. A volume of 5 ml was pressed two times through the filter. The filter was washed with Milli-Q water (Merck Millipore Ltd, Tullagreen, Cork, Ireland) and then dehydrated in ethanol (gradually increasing concentration). Dehydrated samples were placed in microporous capsules of 30 μm in pore size (Ted Pella Inc.) and immersed in absolute ethanol. Critical point drying was performed in an Autosamdri 814 (Tousimis). Once dried, samples were placed on SEM stubs by means of silver conducting paint TAAB S269. Stubs were examined under a scanning electron microscope Hitachi S-4100.

### 2.3. 16S-rDNA analyses with Ion Torrent

DNA was retrieved from sauna and dishwasher samples using the PowerSoil DNA Isolation Kit (MO BIO Laboratories, USA). DNA quality was analyzed using a Nanodrop-1000 Spectrophotometer (Thermo Scientific, Wilmington, DE, USA). A 500 bp long fragment from the hypervariable 16S-rDNA regions V1 – V3 was amplified using the universal primers 28F (5′ -GAG TTT GAT CNT GGC TCA G-3′) and 519R (5′ -GTN TTA CNG CGG CKG CTG-3′). The quality of the resulting amplicons was checked on a 0,8% (w/v) agarose gel. Amplicons were precipitated with 3M potassium acetate and isopropanol. Sequencing libraries were constructed using 100 ng of the DNA pool and performing the amplicon fusion method (Ion Plus Fragment Library Kit, MAN0006846, Life Technologies). Both libraries (Sauna and Dishwasher) were quantified with the Agilent2100 Bioanalyzer (Agilent Technologies Inc, Palo Alto, CA, USA) prior to clonal amplification. Emulsion PCRs were carried out with the Ion PGM Template OT2 400 kit as described following the user guide provided by the manufacturer (MAN0007218, Revision 3.0 Life Technologies). Libraries were sequenced in an Ion 318 Chip v2 on a Personal Genome Machine (PGM) (IonTorrentTM, Life Technologies) at Life Sequencing S.L. (Life Sequencing,Valencia, Spain), using the Ion PGM Sequencing 400 kit and following the manufacturer’s protocol (publication number MAN0007242, revision 2.0, Life Technologies). Short reads (<100bp) and low quality reads (<q15) were removed upon sequencing at the sequencing center. Resulting sequences were analyzed by phylotyping with the MOTHUR software [16]. Amplicons were aligned to the 16S-reference from the Greengenes database. Classification was performed using the k-mer algorithm. Assignments with a similarity percentage lower than 80% were discarded.

### 2.4. Isolation of microbial strains

Lysogenic broth (LB) and Reasoner’s 2A (R2A) agar [17] media were used for bacterial culturing. Samples were suspended in PBS-buffer, vortexed, spread on LB and R2A plates and incubated at 37 °C and 55 °C for one day. Thermophilic and thermoresistant strains were picked, grown in liquid culture and stored in 20 % Glycerol at -70 °C.

### 2.5. Lipolytic Activity and microbial identification

Tributyrin-containing medium is frequently used when screening for lipase-producing microorganisms [18,19], as the degradation of this compound generates clear halos around the lypolitic colonies in the otherwise turbid medium. Samples (1 μL) from *the c*ryo-preserved strains were directly spotted on minimal medium [20], which contained tributyrin (10 mL/L) as main carbon source. Incubations were performed at 4 °C, 20 °C, 37 °C, 46 °C and 55 °C. After 5 days of incubation, the diameter of the halos around lipase-producing strains was measured.

### 2.6. 16S rRNA sequencing of selected strains

Hypervariable 16S-rDNA regions V1 – V3 of the selected strains were amplified by colony PCR using 28F and 519R primers and sequenced with the Sanger method by the Sequencing Service of the University of Valencia (Spain). This allowed the identification of the five selected isolates at a genus level. In order to identify the isolates at a species level, further *Bacillus* spp. primers were used to amplify: the TU elongation factor (tufGPF and tufGPR) [21], a group-specific 16S rRNA region (B-K1/F and B-K1/R1) [22], an endoglucanase gene (ENIF and EN1R) [23] and a glycolsyltransferase (Ba-G206F and Ba-G1013R) [24]. The resulting sequences were manually edited using Pregap4 (Staden Package, 2002) to eliminate low-quality base calls. The final sequence for each isolate was compared to sequence databases using the NCBI BLAST tool.

### 2.7. Lipolytic assays varying pH and salt conditions

Lipase production of the five selected strains was tested on solid minimal medium supplemented with tributyrin (10 mL/L), adjusted to a range of pH (6.5, 8, 9.5 and 11.5), and with or without additional 4 % NaCl. Two microliters of each strain were spotted on each combination of pH and salt media and incubated at 4, 20, 37, 46 or 55 °C for 5 days. After incubation, the diameters of the halos were measured.

In order to determine the optimal conditions for the lipase production of the five selected strains, two microliters of each strain were spotted on additional combinations of pH and salt (pH 6.5, 8, 9.5 and 11.5; NaCl 0, 1, 2, 3 and 4 %). The plates were incubated for five days at 37 °C. The assay was performed in triplicate.

## 3. Results and Discussion

### 3.1. Scanning Electron Microscopy

The samples obtained from a wet sauna and a dishwasher filter proved rich in microorganisms, as deduced by observation under SEM (Fig 1). In the sauna sample, microorganisms were mostly present in the form of a very dense biofilm almost totally embedded in a smooth matrix, very likely made of EPS (Fig 1A); whereas the dishwasher filter sample consisted mainly of food debris with scattered microorganisms (Fig 1B).

**Fig 1.**
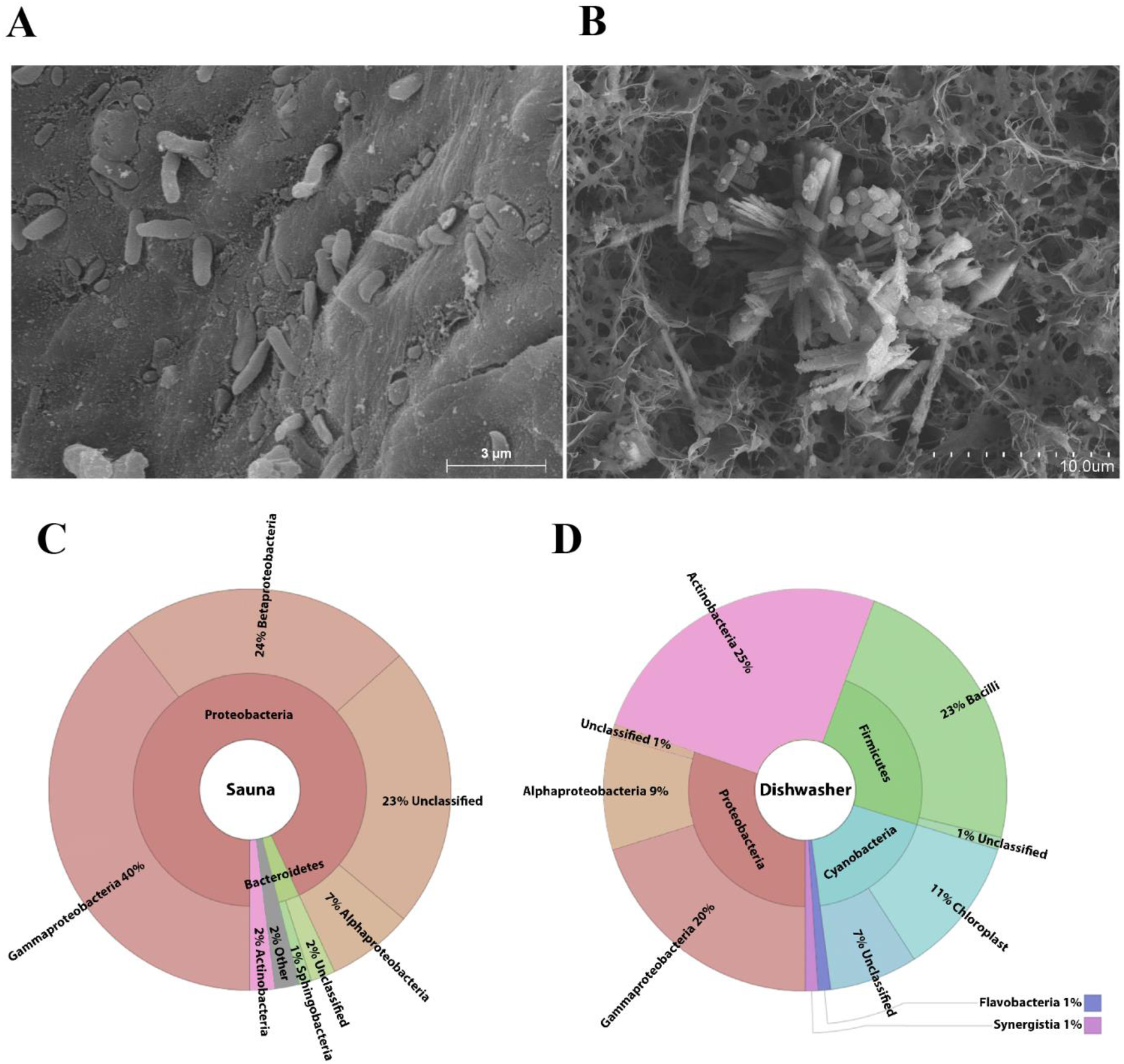
Scanning electron micrographs from the sauna. (A) and the dishwasher (B) samples; taxonomic diversity estimated by 16Sr amplicon sequencing of sauna (C) and dishwasher (D). The sauna (C) sample was especially rich in Proteobacteria; whereas the dishwasher filter (D) also contained high amounts of Firmicutes, Cyanobacteria and Actinobacteria.

### 3.2. 16S-rDNA analyses with Ion Torrent

The taxonomic diversity of the two samples was determined by high throughput-sequencing, performed as described in Materials and Methods, and resulted in very different taxonomic profiles of both samples (Fig 1C and 1D). Proteobacteria were overwhelmingly abundant in the sauna sample, accounting for more than 90 % of the reads (Fig 1C). Of those, alpha-, beta- and gamma-proteobacteria were present at similar frequencies, each accounting for more than 20 % of the assigned sequences. Minor taxa with frequencies of 1-5% included Bacteroidetes, Actinobacteria and Acidobacteria. The dishwasher filter was characterized by large amounts of Proteobacteria, Firmicutes (Bacilli, most of them), Cyanobacteria and Actinobacteria; and very low amounts of other taxa (Fig 1D).

These results are consistent with previous reports on these two extreme environments. Lee *et al.* [25] characterized the bacterial community contaminating the floor of a hot and dry sauna, which proved rich in Firmicutes, Gamma-proteobacteria and Beta-proteobacteria. Another report by Kim *et al.* [26] of a 64°C dry sauna revealed a population with Firmicutes, Gamma-proteobacteria, Beta-proteobacteria and Deinococci as the most frequent taxa. As mentioned above, our samples were rich Beta- and Gammaproteobacteria, although we also detected Alpha-proteobacteria, which was absent in the works by Kim *et al.* [25] and Lee *et al*. [26]. Reciprocally, we did not detect Firmicutes or Deinococci with our 16S rRNA analysis, while both taxa were found by those two previous reports. Concerning the dishwasher samples, a previous report by Savage *et al.* [27] characterized, among other household surfaces, the bacteria present in the dishwasher rinse reservoir. According to that previous report, bacterial population in the dishwasher consists of Proteobacteria, Firmicutes, Cyanobacteria and Actinobacteria, which corresponds to the taxonomic profile we found in the dishwasher filter. Nevertheless, Euryarcheota and Bacteroidetes that were found in the rinse reservoir [27] were not detected in the filter in the present work.

### 3.3. Culturing strains and lypolitic activity screening

Bacterial colonies randomly selected among those with a strong growth on LB and R2A plates were re-streaked to yield a collection of strains, and lipase production was screened after growth for five days in tributyrin-containing media, as described in materials and methods. Lipase production of all the strains is shown in Fig 2A. Most of the strains (72 and 79 % of the sauna and dishwasher strains, respectively) displayeds some level of lipolytic activity. In general, sauna strains were able to produce lipases within a broader range of temperature, including, in two cases, values as high as 46 °C and as low as 4 °C. At least in these cases, though, lipases are not only produced, but are fully functional at extreme temperatures as deduced by this assay. In contrast, strains from the dishwasher displayed lipolytic activity within a smaller range of temperatures, in most cases only around 37 °C within the tested range. Interestingly, lipolytic activity, as deduced by haloes diameter, was maximum at 46 °C in several sauna strains, although no strains produced detectable lipolysis at 55 °C.

**Fig 2.**
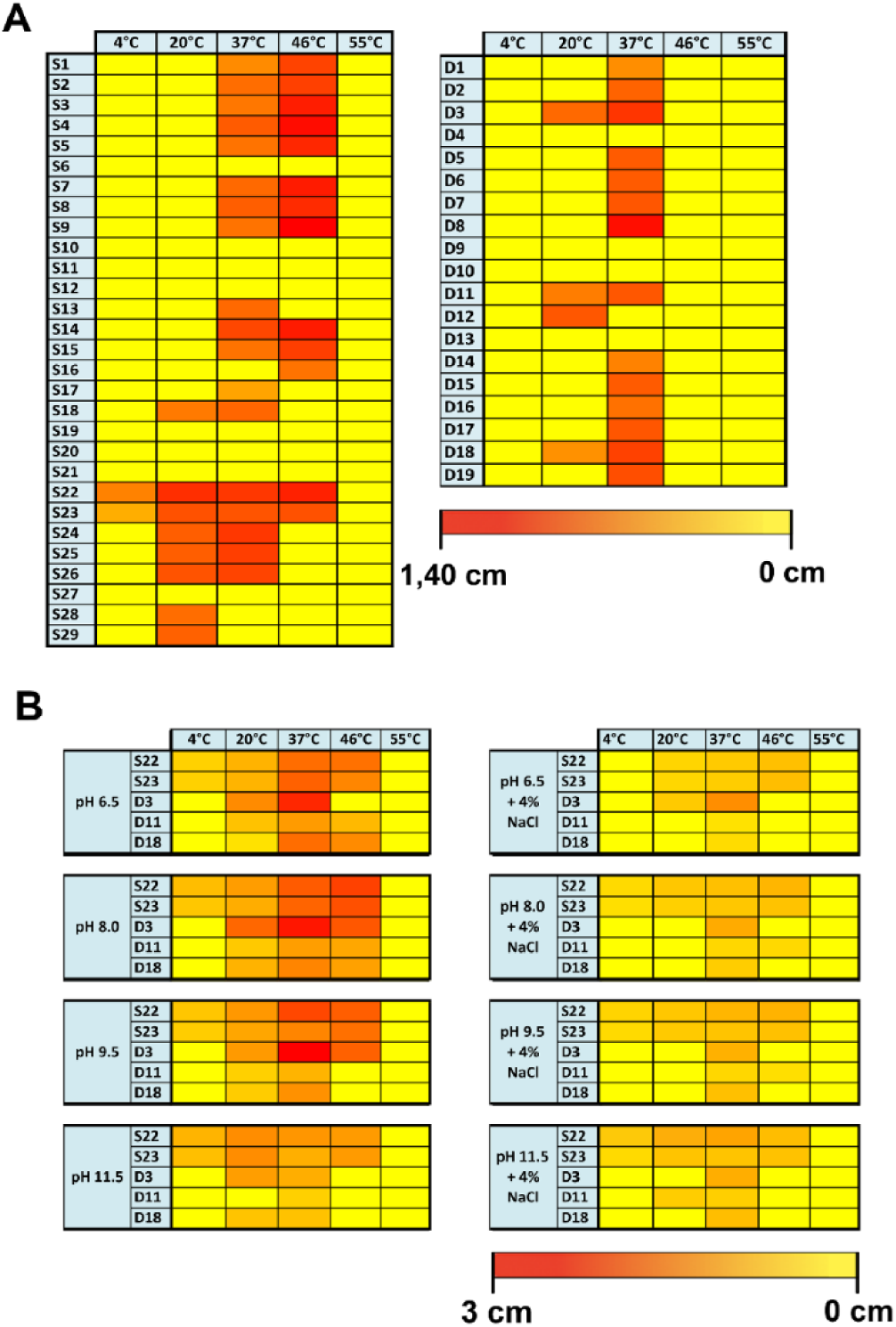
Heatmaps of lipolytic activity. (A) Isolates from the sauna and from the dishwasher contained mainly strains with lipolytic activity. Strains from the sauna exposed lipolytic activity within a broader range of temperature. Sauna samples with the widest range of temperature were S22 and S23. In the dishwasher most samples showed lipolytic activity at 37 °C and only sample D3, D11 and D18 had lipolytic activity at two tested temperatures. (B) Heatmap of lipolytic activity of selected strains from sauna (S22, S23) and dishwasher (D3, D11, D18) under different pH (6.5, 8, 9.5, 11.5) and temperature conditions (4, 20, 37, 46 and 55°C) in minimal medium or minimal medium with 4 % NaCl.

On the basis of this first screening for lypolitic activities, five strains were selected for further assays. Those included the strains with the broadest temperature activity range: S22 and S23, both active in all temperatures tested except at 55 °C; and D3, D11 and D18, the three dishwasher samples active at both 20 °C and 37 °C. In order to assess the potential of these five strains for biotechnological purposes (in terms of lipase production under extreme environmental conditions), they were taxonomically identified and subjected to a stress test under a range of temperatures and pH conditions, performed in minimum media with and without 4% added salt. Lipase activity were tested under different temperatures, NaCl and pH conditions, and results are shown in Fig 2B. Again, sauna samples exhibited a broad range of thermal stability, with medium to large halos at pH values mildly acid to moderately alkaline (6.5-9.5) and even in the presence of 4 % NaCl (pH 8 and 9.5). Interestingly, very alkaline (11.5) conditions, combined with high salt contents correlated with an increased thermal range of lipase production and activity for both S22 and S23, which increased from 20 °C up to 46 °C. In general, though, salt addition yielded smaller haloes at any temperature compared to standard media.

Regarding the strains we isolated from the dishwasher filter, assays performed with minimum media (without salt) adjusted to a wide range of pH values and incubated at different temperatures revealed the alkaliphility of their lipolytic abilities, both in terms of thermal broad range at alkaline pH values, and intensity of the activity as deduced by haloes sizes (Fig 2B). Addition of NaCl to the media resulted in smaller haloes and, at least for pH values of 8-9.5, in a narrower thermal activity range. In the three dishwasher strains, the combination of 4 % added NaCl and high (11.5) pH resulted in an altered thermal range of activity. At least in one case (D11) addition of 4 % NaCl partially restored the lack of activity observed with no added salt and at a pH of 11.5.

Sequencing of a16S rRNA gene fragment allowed identification of all five isolates as *Bacillus sp.* Further sequencing of the TU elongation factor (tufGP primers) and the group-specific 16S rRNA region (BK-1 primers) revealed D11 strain as *Bacillus megaterium* (with 99 and 100 % identity, respectively); and S22/S23 strains as *Bacillus pumilus* (with 100 and 99 % identity in the case of tufGP and BK-1 primers, respectively). D3 was not completely identified, and remains as *Bacillus* sp., possibly *B. subtilis, B. amyloliquefaciens, B. methylotrophicus* or *B. velenznesis*. D18 was identified as *B. cereus/B. thuringiensis.*

### 3.4. Robustness of the selected isolates in varying pH and salt conditions

In order to characterize the robustness of lypolitic activity of the five selected strains under alkaline and/or high salinity conditions, the strains were tested on combined pH and salt contents conditions, at 37 °C. Lypolitic activity results are shown in Fig 3.

**Fig 3.**
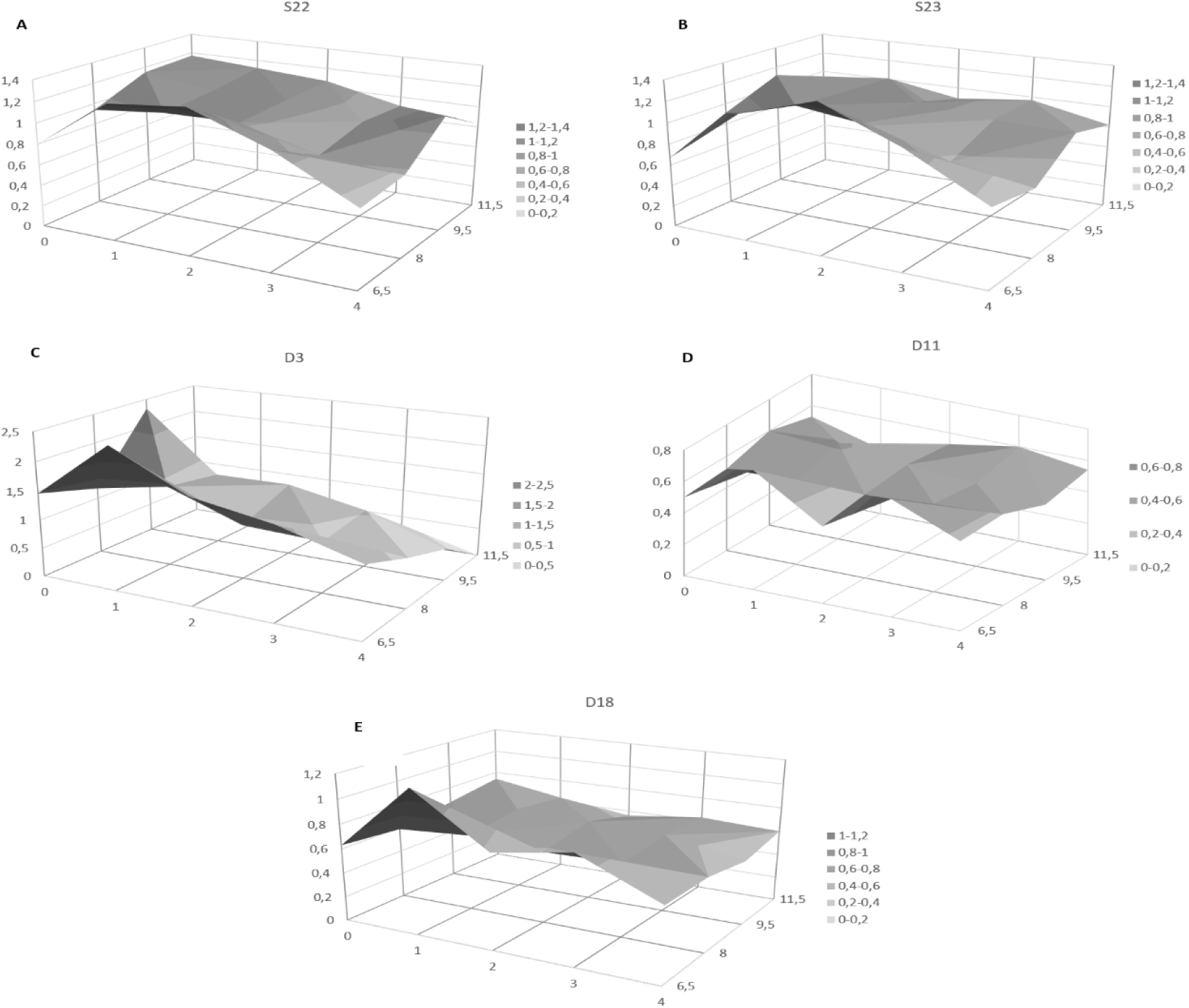
Surface graphs of lypolitic activities of the five selected strains (S22, S23, D3, D11 and D18) under different pH (6.5, 8, 9.5 and 11.5) and NaCl (0, 1, 2, 3 and 4%) conditions. Diameter of the lipolysis haloes (cm) is represented in the Y axis, whereas salt and pH conditions are represented in the X and Z axis, respectively. Halo sizes are in grey to dark grey as indicated at the bottom of each graph.

In general, all strains displayed a very robust lipase activity under different salt and pH conditions when grown at 37 °C, as deduced by the relatively flat 3D profile (Fig 3), although halo diameters generally decreased towards extreme salt values (0% and 4% NaCl). Specifically, D11 and D18 (Fig 3D and 3E, respectively) were the most robust lipase producers, followed by S22 and S23 (Fig 3A and 3B, respectively). In contrast, D3 (Fig 3C) was the least robust strain, with variations in activity depending on pH conditions and an even higher salt-dependent variation: lipolytic activity ranged from undetectable in very alkaline conditions to very large (2.45 cm) haloes at pH 6.5 with 1% NaCl.

Aside from the robustness observed, the five selected strains displayed clear optimum peaks at pH 6.5 with 2% NaCl for S22 and S23; and with 1% NaCl for D3, D11 and D18. D3 displayed the highest lypolitic activity under optimum conditions, with halo diameters of up to 2.45 cm, followed by S22 and S23, both with diameters of up to 1.35 and 1.40 cm, respectively. Strains D11 and D18 displayed the lowest lipolytic activity, with maximum halos of 0.76 and 1.17, respectively.

*Bacillus* sp. have been previously reported to produce thermostable lipases [28, 29, 30, 31]. Regarding the *Bacillus* species that we have isolated and identified in the present work, they are known to produce thermo-resistant lipases, some of them stable at very low or very high pH values. For example, thermostable lipases can be found in *B. megaterium* (a monoacylglycerol lipase and a carboxylesterase, Uniprot accession number: A0A0H4RCB5 and G2RXU5). Furthermore, a thermostable extracellular lipase has been described for this species, which is capable of retaining 100 % of its activity at 50 °C, and becomes stimulated in the presence of acetone, DMSO, isopropanol and several reducing agents [32]. On the other hand, there are several thermostable lipases known for the *B. cereus* group (Uniprot accession number: A0A0B5NXJ9; A0A090YL00; A0A0A0WM49) and for the *B. subtilis* group [33, 34]. *Bacillus pumilus* is well-known for its thermostable lipases, as many reports describe lipase fully or partially functional at high temperatures [35, 36, 37, 38, 39], including a lipase that is able to resist temperatures up to 100 °C [36]. Even lipases that are functional at both high temperatures and high or low pH values have been described [37, 38]. Finally, *B. pumilus*, has been identified as the most efficient lypolitic enzymes producer out of 65 strains analysed in a previous report [39]. In fact, in our experiments, isolates S22 and S23 were among the three strains with the highest lypolitic activity, and both of these were identified as *B. pumilus.*

In summary, we have identified from domestic environments several *Bacillus* spp. which are strong producers of robust lypolitic enzymes, and this is in concordance with the literature on strains of this genus isolated from other environments. It has to be highlighted that, in our assays, the lipases corresponding to the isolated strains showed high activity at wide ranges of pH (6.5 – 11.5), temperature (4°C - 46 °C) and salt (up to 4 % NaCl), and that all measurements were performed in-situ within their host organisms. Therefore, not only the bacteria are able to produce lipases under these conditions, but also these lipases are perfectly functional. Further tests of the lipase extracts will shed light on the robustness of the lipolytic activity itself, without the limitations caused by the bacterial production.

Our results suggest that the isolated strains may be used as robust chassis for lipase production, and these lipases may be used in the industry as robust bio-detergents. As described in the works mentioned above, especially *B. pumilus* seems to be of interest, since it shows a very strong lipolytic activity and since it adapts, according to our data, most efficiently to different conditions of pH, salt and temperature. This is especially interesting, since we found no previous description of *Bacillus* lipases, which work efficient in wide range of pH, salt and temperature at the same time. The present work shows for the first time the potential of domestic environments as a source of *Bacillus* strains with potential biotechnological applications.

## 4. Conclusions

The present work is the first screening of extreme indoor environments specifically aiming at the identification of biotechnological relevant bacterial strains able to produce robust enzymes, in our case, robust lipases. Our results reveal that such domestic environments are promising sources for the identification of robust enzymatic activities, as we have managed to isolate five strains with stable lipolytic activity under a wide range of temperature, salt and pH conditions. These metabolic capabilities can be especially useful as components of robust bio-detergents. Furthermore, this work might be the first step of a new view on the human-associated indoor microbiome, focused on ecological aspects and on biotechnological applications.

## Acknowledgements

We thank C. Vilanova and X. Baixeras for assistance with electron microscopy. Further the authors are indebted to E. L. James for inspiring us to choose the title for the present work.

---

**S1 Data. Sequences obtained from high-throughput 16S rRNA sequencing of the DNA isolated from the sauna sam-ple.**

**S2 Data. Sequences obtained from high-throughput 16S rRNA sequencing of the DNA isolated from the dishwasher sample.**

**Both available under request.**

